# MicroPheno: Predicting environments and host phenotypes from 16S rRNA gene sequencing using a k-mer based representation of shallow sub-samples

**DOI:** 10.1101/255018

**Authors:** Ehsaneddin Asgari, Kiavash Garakani, Alice C. McHardy, Mohammad R.K. Mofrad

## Abstract

**Motivation:** Microbial communities play important roles in the function and maintenance of various biosystems, ranging from the human body to the environment. A major challenge in microbiome research is the classification of microbial communities of different environments or host phenotypes. The most common and cost-effective approach for such studies to date is 16S rRNA gene sequencing. Recent falls in sequencing costs have increased the demand for simple, efficient, and accurate methods for rapid detection or diagnosis with proved applications in medicine, agriculture, and forensic science. We describe a reference- and alignment-free approach for predicting environments and host phenotypes from 16S rRNA gene sequencing based on k-mer representations that benefits from a bootstrapping framework for investigating the sufficiency of shallow sub-samples. Deep learning methods as well as classical approaches were explored for predicting environments and host phenotypes.

**Results:** k-mer distribution of shallow sub-samples outperformed the computationally costly Operational Taxonomic Unit (OTU) features in the tasks of body-site identification and Crohn's disease prediction. Aside from being more accurate, using k-mer features in shallow sub-samples allows (i) skipping computationally costly sequence alignments required in OTU-picking, and (ii) provided a proof of concept for the sufficiency of shallow and short-length 16S rRNA sequencing for phenotype prediction. In addition, k-mer features predicted representative 16S rRNA gene sequences of 18 ecological environments, and 5 organismal environments with high macro-F1 scores of 0.88 and 0.87. For large datasets, deep learning outperformed classical methods such as Random Forest and SVM.

**Availability:** The software and datasets are available at https://llp.berkeley.edu/micropheno.

## 1 Introduction

Microbial communities have important functions relevant to supporting, regulating, and in some cases causing unwanted conditions (e.g. diseases) in their hosts/environments, ranging from organismal environments, like the human body, to ecological environments, like soil and water. These communities typically consist of a variety of microorganisms, including eukaryotes, archaea, bacteria and viruses. The human microbiome is the set of all microorganisms that live in close association with the human body. It is now widely believed that changes in the composition of our microbiomes correlate with numerous disease states, raising the possibility that manipulation of these communities may be used to treat diseases. Normally, the microbiota (particularly the intestinal microbiota) of the microbiome play several important roles in humans, which include: (i) prevention of pathogen growth, (ii) education and regulation of the host immune system, and (iii) providing energy substrates to the host [1]. Consequently, dysbiosis of the human microbiome can promote several diseases, including asthma [2, 3], irritable bowel syndrome [4, 5], Clostridium difficile infection [6], chronic periodontitis [7, 8], cutaneous leishmaniasis [9], obesity [10, 11], chronic kidney disease [12], Ulcerative colitis [13], and Crohns disease [14, 15]. The human microbiome appears to play a particularly important role in the development of Crohns disease. Crohns disease is an inflammatory bowel disease (IBD) with a prevalence of approximately 40 per 100,000 and 200 per 100,000 in children and adults, respectively [16].

Microbiomes present in the environment also serve important functions, such as nutrient cycling [17]. Due to differences in nutrient availability and environmental conditions, microbiomes in different environments are each characterized by unique structures and microbial compositions [18–21]. Furthermore, each microbiome plays a unique role in the environment. For instance, the ocean microbiome generates half of the primary production on Earth [19]. The soil microbiome surrounding the root of plants has a great impact on plant fertility and growth [22]. Additionally, analyzing microbial communities in drinking water is one of the primary concerns of the environmental sciences [18].

The starting point in data collection for many of the above mentioned projects is 16S rRNA gene sequencing of microbial samples [23]. There are a number of features that make the 16S rRNA gene ideal for use as a taxonomic ‘fingerprint’ for microbiome composition characterization. First, the 16S rRNA gene is highly conserved across bacteria and archaea. Secondly, the gene consists of both conserved regions, for which universal species-independent PCR primers may be directed against, and nine hypervariable regions (V1-V9), along which species-specific sequence differences may accumulate to allow for differential identification of species [24]. Sequencing of specific hypervariable regions of the 16S rRNA gene may be performed for determination of microbial community composition [25]. After the 16S rRNA hypervariable regions are sequenced from a microbial sample, the obtained sequences are then processed using bioinformatics software (such as QIIME [26, 27], Mothur [28], or Usearch [29]]) and clustered into groups of closely related sequences referred to as Operational Taxonomic Units (OTUs), which may then be used to assist in the functional profiling of microbial samples. Later inx1.2 we discuss the pros and cons of OTU features in details.

### 1.1 Machine learning for environments or host phenotypes classification

Several recent studies predicted the environment or host phenotypes using 16S gene sequencing data for body-sites [30, 31], disease state [32–35], ecological environment quality status prediction [36], and subject prediction for forensic science [37, 38]. In all, OTUs served as the main input feature for the down stream machine learning algorithms. Random Forest and then, ranking second, linear Support Vector Machine (SVM) classifiers were reported as the most effective classification approaches in these studies [31, 39, 34].

Related prior work on body-site classification [30, 31] used the following datasets: Costello Body Habitat (CBH - 6 classes), Costello Skin Sites (CSS - 12 classes) [40], and Pei Body Site (PBS - 4 classes)[31]. An extensive comparison of classifiers for body-site classification over CBH, CSS, and PBS on top of OTU features has been performed by Statnikov et al [31]. The best accuracy levels measured by relative classifier information (RCI) achieved by using OTU features are reported as 0.784, 0.681, and 0.647 for CBH, CSS, and RCI respectively. Due to the insufficiency of the number of samples (on average 57 samples per class for CBH, CSS, and PBS) as well as the unavailability of raw sequences for some of the datasets mentioned above, instead of using the same dataset we replicate the state-of-the-art approach suggested in [31], i.e. Random Forest and SVM over OTU features for a larger dataset (Human Microbiome Project dataset). We then compare OTU features with k-mer representations. Working on a larger dataset allows for a more meaningful investigation and better training for deep learning approaches.

Detecting disease status based on 16S gene sequencing is becoming more and more popular, with applications in the prediction of Type 2 Diabetes [35] (patients: 53 samples, healthy:43 - Best accuracy: 0.67), Psoriasis (151 samples for 3 classes - Best accuracy: 0.225), IBD (patients: 49 samples, healthy:59 - Best AUC:0.95) [32], (patients: 91 samples, healthy: 58 samples - Best AUC:0.92) [33]. Similar to body-site classification datasets, the datasets used for disease prediction were also relatively small. In this paper, we use the Crohn’s disease dataset [14] with 1359 samples (patients: 731 samples, negative class: 628 samples) for evaluating our proposed method and then compare it with the use of OTU features.

We focus on machine learning approaches for classification of environments or host phenotypes of 16S rRNA gene sequencing data, which is the most popular and cost-effective sequencing method for the characterization of microbiome to date [39]. Studies on the use of machine learning for predicting microbial phenotype instead of environments/host phenotype [41, 42], as well as predictions based on shotgun metagenomics and whole-genome microbial sequencing are beyond the scope of this paper, although we believe that one may easily adapt the proposed approach to shotgun metagenomics, similar to the study by Cui et al. on IBD prediction [43].

Recently, deep learning methods became popular in various applications of machine learning in bioinformatics [44, 45] and in particular in metagenomics [46]. However, to the best of our knowledge this paper is the first study exploring environment and host phenotype prediction from 16S rRNA gene sequencing data using deep learning approaches.

### 1.2 16S rRNA gene sequence representations

#### OTU representation

As reviewed inx1.1, prior machine learning works on environment/host phenotype prediction have been mainly using OTU representations as the input features to the learning algorithm. Almost all popular 16S rRNA gene sequence processing pipelines cluster sequences into OTUs based on their sequence similarities utilizing a variety of algorithms [47, 27]. QIIME allows OTU-picking using three different strategies: **(i) closed-reference OTU-picking:** sequences are compared against a marker gene database (e.g. Green-genes [48] or SILVA [49]) to be clustered into OTUs and then the sequences different from the reference genomes beyond a certain sequence identity threshold are discarded. (ii) open-reference OTU-picking: the remaining sequences after a closed-reference calling go through a de novo clustering. This allows for using the whole sequences as well as capturing sequences belonging to new communities which are absent in the reference databases [50]. (iii) pure de novo OTU-picking: sequences (or reads) are only compared among themselves and no reference database is used. The third strategy is more appropriate for novel species absent in the current reference. Although OTU clustering reduces the analysis of millions of reads to working with only thousands of OTUs and simplifies the subsequent phylogeny estimation and multiple sequence alignment, OTU representations have several shortcomings: (i) All three OTU-picking strategies involve massive amounts of sequence alignments either to the reference genomes (in closed/opened-reference strategies) or to the sequences present in the sample (in open-reference and de novo strategies) which makes them very expensive [51] in comparison with reference-free/alignment-free representations. **(ii)** Overall sequence similarity is not a proper condition for grouping sequences and OTUs can be phylogenetically incoherent. For instance, a single mutation between two sequences is mostly ignored by OTU-picking algorithms. However, if the mutation does not occur within the sample, it might be a signal for assigning a new group. In addition, several mutations within a group most likely are not going to be tolerated by OTU-picking algorithms. However, having the same ratio across samples may suggest that the mutated sequences belong to the same group [52, 47]. (iii) The number of OTUs and even their contents are very sensitive to the pipeline and parameters, and this makes them difficult to reproduce [53].

#### k-mer representations

k-mer count vectors have been shown to be successful input features for performing machine learning on biological sequences for a variety of bioinformatics tasks [54]. In particular, recently k-mer count features have been largely used for taxonomic classifications of microbial metagenomics sequences [55–60]. However, to the best of our knowledge, k-mer features have not been explored for phenotypical and environmental characterizations of 16S rRNA gene sequencing. The advantages of using k-mer features over OTUs are later discussed in the discussion section.

In this paper we propose using shallow sub-samples for predicting environments and host phenotypes from 16S rRNA sequencing. Our approach is fast, reference-free and alignment-free, while contributing in building accurate classifiers outperforming conventional OTU features in body-site identification and Crohn’s disease classification. We propose a bootstrapping framework to investigate the sufficiency of shallow sub-samples for the prediction of the phenotype of interest, which proves the sufficiency of short-length and shallow sequencing of 16S rRNA. In addition, we explore deep learning methods as well as classical approaches for the classification and show that for large datasets, deep learning can outperform classical methods. Furthermore, we explore PCA, t-SNE, and supervised deep representation learning for visualization of microbial samples/sequences of different phenotypes. We also show that k-mer features can be used to predict representative 16S rRNA gene sequences belonging to 18 ecological environments and 5 organismal environments with high macro-F1s.

## 2 Material and Methods

### 2.1 Datasets

#### Body-site identification

We employ the metagenomic 16S rRNA gene sequence dataset provided by the NIH Human Microbiome Project (HMP) [61, 62]^1^. In particular, we use processed, annotated 16S rRNA gene sequences of up to 300 healthy individuals, each sampled at 4 major body-sites (oral, airways, gut, vagina) and up to three time points. For each major body-site, a number of sub-sites were sampled. We focus on 5 body sub-sites: anterior nares (nasal) with 295 samples, saliva (oral) with 299 samples, stool (gut) with 325 samples, posterior fornix (urogenital) with 136 samples, and mid vagina (urogenital) with 137 samples, in total 1192 samples. These body-sites are selected to represent differing levels of spatial and biological proximity to one another, based on relevance to pertinent human health conditions potentially influenced by the human microbiome. In order to compare k-mer based approach with state-of-the-art OTU features we collect the closed-reference OTU representations of the same samples in HMP [62] ^2^ obtained using the QIIME pipeline [50].

#### Crohn’s disease prediction

For the classification of Crohns disease, we use the metagenomics 16S rRNA gene sequence dataset described in [14]^3^, which is currently the largest pediatric Crohns disease dataset available. This dataset included annotated 16S rRNA gene sequence data for 731 pediatric (≤ 17 years old) patients with Crohns disease and 628 samples verified as healthy or diagnosed with other diseases, making a total of 1359 samples. The 16S rRNA sequencing dataset was targeted towards the V4 hypervariable region of the 16S rRNA gene. Similar to the body-site dataset, in order to compare the k-mer based approach with the approach based on OTU features we collect the OTU representations of the same samples from Qiita repository ^4^ obtained using QIIME pipeline [50].

#### Prediction of the environment for representative 16S rRNA gene sequences

MetaMetaDB provides a comprehensive dataset of representative 16S rRNA gene sequences of various ecological and organismal environments collected from existing 16S rRNA databases spanning almost 181 million raw sequences. In the MetaMetaDB pipeline, low-quality nucleotides, adaptors, ambiguous sequences, homopolymers, duplicates, and reads shorter than 200bp, as well as chimera have been removed and finally 16S rRNA sequences are clustered with 97% identity generating 1,241,213 representative 16S rRNA sequences marked by their environment [63]. MetaMetaDB divides its ecological environments into 34 categories and its organismal environments into 28 categories. We create three datasets which are subsets of MetaMetaDB to investigate the discriminative power of k-mers in predicting microbial habitability. Since the sequences in MetaMetaDB were already filtered and semi-identical sequences were removed, OTU-picking would not be relevant as it would result in an almost one-to-one mapping between the sequences and OTUs (we verified this using QIIME).

##### Ecological environment prediction

MetaMetaDB is imbalanced in terms of the number of representative sequences per environment. For this study, we pick the ecological environments with more than 10,000 samples, ending up with 18 classes of ecological environments: activated sludge, ant fungus garden, aquatic, bioreactor, bioreactor sludge, compost, food, food fermentation, freshwater, freshwater sediment, groundwater, hot springs, hydrocarbon, marine, marine sediment, rhizosphere, sediment, and soil ^5^. We make two datasets out of the sequences in these environments: ECO-18K containing 1000 randomly selected instances per class (a total of 18K sequences) and ECO-180K, which is 10 times larger than ECO-18K, i.e. contains 10,000 randomly selected instances per class (a total of 180K sequences).

**Organismal environment prediction** from the organismal environments in MetaMetaDB we select a subset containing gut microbiomes of 5 different organisms (bovine gut, chicken gut, human gut, mouse gut, termite gut) and down-sample each class to the size of the smallest class ending up having 620 sample per class (in total containing 3100 sequences) we call this dataset 5GUTS-3100.

### 2.2 MicroPheno computational workflow

We describe using deep learning and classical methods for classification of the environments or host phenotypes of microbial communities using k-mer frequency representations obtained from shallow sub-sampling of 16S rRNA gene sequences. We propose a bootstrapping framework to confirm the sufficiency of using a small portion of the sequences within a 16S rRNA sample for determining the underlying phenotype.

The MicroPheno computational workflow has the following steps (Figure 1): (i) to find the proper size *N* for the sample, such that it stays representative of the data and produces a stable k-mer profile, the 16S rRNA sequences go through a bootstrapping phase. (ii) Afterwards, the sub-sampled sequences are used to find the best value for k for classification, to produce the k-mer representations of the samples. (iii) The k-mer representations are used for classification with Deep Neural Networks (DNN), Random Forest (RF), and Linear SVM. (iv) Finally, the k-mer representations as well as the supervised representations trained using DNNs are used for visualization of the 16S rRNA gene sequences or samples. In what follows, these steps are explained in detail.

**Figure 1:**
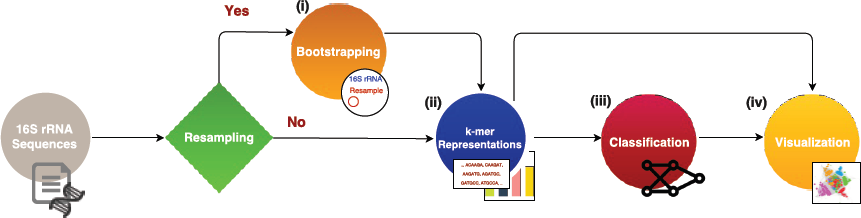
The components and the data flow in the MicroPheno computational workflow

#### Bootstrapping

Confirming the sufficiency of only a small portion of 16S rRNA sequences for environment/phenotype classification is important because (i) sub-sampling reduces the preprocessing run-time, and (ii) more importantly, it proves that even a shallow 16S rRNA sequencing is enough. For this purpose we propose a resampling framework similar to bootstrapping to give us quantitative measures for finding the proper sampling size. Let *θ_k_*(*X_i_*) be the normalized k-mer distribution of *X_i_*, a set of sequences in the *i^th^* 16S rRNA sample. We investigate whether only a portion of *X_i_*, which we represent as 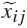, i.e. *j^th^* resample of *X_i_* with sample size N, would be sufficient for producing a proper representation of *X_i_*. To quantitatively find a sufficient sample size for *X_i_* we propose the following criteria in a resampling scheme. (i) **Self-consistency**: resamples for a given size N from *X_i_* produce consistent 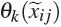’s, i.e. resamples should have similar representations. **(ii) Representativeness**: resamples for a given size N from *X_i_* produce 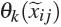’s similar to θ_*k*_(*X_i_*), i.e. similar to the case where all sequences are used.

We quantitatively define self-inconsistency and unrepresentativeness and seek parameter values that minimize them. We measure the **self-inconsistency** 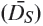 of the resamples’ representations by calculating the average Kullback Leibler divergence among normalized k-mer distributions for *N_R_* resamples (here *N_R_*=10) with sequences of size N from the *i^th^* 16S rRNA sample:

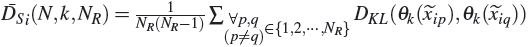

where 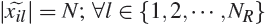. We caculate the average of the values of 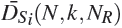 over the *M* different 16S rRNA samples:

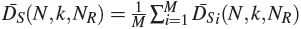

We measure the **unrepresentativeness** 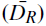 of the resamples by calculating the average Kullback Leibler divergence between normalized k-mer distributions for *N_R_* resamples (*N_R_*=10) with size N and using all the sequences in *X_i_* for the *i^th^* 16S rRNA sample:

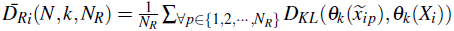

where 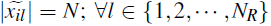. We calculate the average over 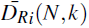’s for the *M* 16S rRNA samples:

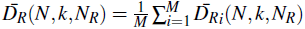

For the experiments on body-site and the dataset for Crohn’s disease, we measure self-inconsistency 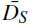 and unrepresentativeness 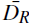 for *N_R_* = 10 and *M* = 10 for any 8 ≥ *k* ≥ 3 with sampling sizes ranging from 20 to 10000. Each point in Figure 5 represents the average of 100 (*M* × *N_R_*) resamples belonging to *M* randomly selected 16S rRNA samples, each of which is resampled *N_R_* = 10 times. Since in the ecological and organismal datasets each sample is a single sequence, the bootstrapping step is skipped.

#### k-mer representation

We propose *l*1 normalized k-mer distribution of 16S rRNA gene sequences as input features for the downstream machine learning classification algorithms as well as visualization. Normalizing the representation allows for having a consistent representation even when the sampling size is changed. For each k-value we pick a sampling size that gives us a self-consistent and representative representation measured by 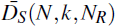 and 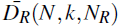 respectively as explained above. MicroPheno provides a parallel python implementation of k-mer distribution generation for a given sampling size.

#### Classification

Random Forests and linear SVM are the state-of-the-art classical approaches for categorical prediction on 16S rRNA sequencing [31, 39, 34] and in general for many machine learning problems in bioinformatics [64]. These two approaches, which are respectively instances of non-linear and linear classifiers, are both adopted in this study. In addition to these classical approaches, we also evaluate the performance of deep Neural Network classifiers in predicting environments and host phenotypes.

We evaluate and tune the model parameter in a stratified 10 fold cross-validation scheme. To ensure optimizing for both precision and recall, we optimize the classifiers for the harmonic mean of precision and recall, i.e. F1. In particular, to give equal importance to the classification categories, specifically when we have imbalanced classes, we use macro-F1, which is the average of F1’s over categories. Finally the evaluation metrics are averaged over the folds and the standard deviation is also reported.

##### Classical learning algorithms

We use a one-versus-rest strategy for multi-class linear SVM [65] and tune parameter C, the penalty term for regularization. Random Forest [66] classifiers are tuned for (i) the number of decision trees in the ensemble, (ii) the number of features for computing the best node split, and (iii) the function to measure the quality of a split.

##### Deep learning

We use the Multi-Layer-Perceptrons (MLP) Neural Network architecture with several hidden layers using Rectified Linear Unit (ReLU) as the nonlinear activation function. We use softmax activation function at the last layer to produce the probability vector that can be regarded as representing posterior probabilities [67]. To avoid overfitting we perform early stopping and also use dropout at hidden layers [68]. A schematic visualization of our Neural Networks is depicted in Figure 2. Our objective is minimizing the loss, i.e. cross entropy between output and the one-hot vector representation of the target class. The error (the distance between the output and the target) is used to update the network parameters via a Back-propagation algorithm using Adaptive Moment Estimation (Adam) as the optimizer [69]. We start with a single hidden layer and incrementally increase the number of layers with systematic exploration of the number of hidden units and dropout rates to find a proper architecture. We stop adding layers when increasing the number of layers does not result in achieving a higher macro-F1 anymore. In addition, for the visualization of samples as explained later in 2.2 we use the output of the (*n* − 1)^*th*^ hidden layer.

**Figure 2.**
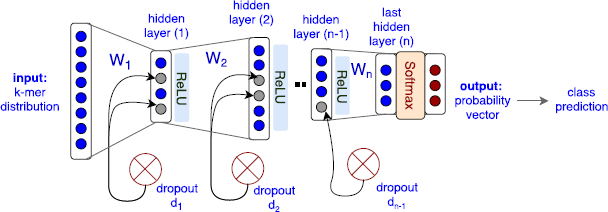
General architecture of the MLP Neural Networks that have been used in this study for multi-class classification of environment and host phenotypes.

#### Visualization

To project 16S rRNA sequencing samples to 2D for visualization purposes, we explore Principal Component Analysis (PCA) [70] as well as t-Distributed Stochastic Neighbor Embedding (t-SNE) [71] as instances of respectively linear and non-linear dimensionality reduction methods. In addition, we explore the use of supervised deep representation learning in visualization of data [72], i.e. we visualize the activation function of the last hidden layer of the Neural Network trained for prediction of environments or host phenotypes to be compared with unsupervised methods. More details on visualization methods are provided as supplementary materials.

##### Implementations

MicroPheno uses implementations of Random Forest, SVM, t-SNE, and PCA in the Python library scikit-learn [73] and Deep Neural Networks are implemented in the Keras^6^ deep learning framework using TensorFlow back-end.

## 3 Results

In this section, the results are organized based on datasets. As discussed in Section 2.2 we have several choices in each step in the computational workflow: choosing the value of k in k-mer, the sampling rate, and the classifiers. To explore the parameter space more systematically we followed the steps demonstrated in Figure 3. (i) In the first step for each value of 8 ≥ *k* ≥ 3 we pick a stable sample size based on the output of bootrapping. (ii) As the next step, we perform the classification task using tuned Random Forest for different k values and their selected sampling sizes based on boostrapping. We selected Random Forest because we found it easy to tune in addition to the fact that it outperforms linear SVM in many cases. (iii) As the third step, for a selected k we investigate the role of sampling size (N) in classification. (iv) Finally, we compare different classifiers for the selected k and N. We also compare the performance of our proposed k-mer features with that of OTU features in classification tasks.

**Figure 3.**
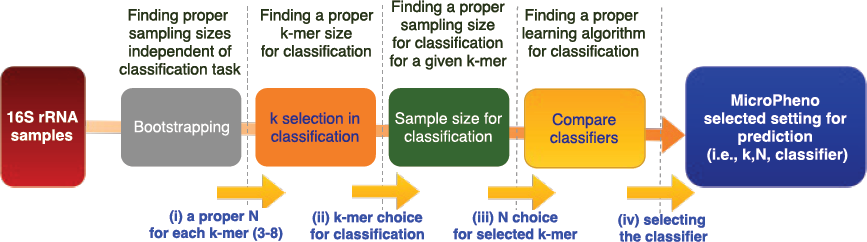
Steps we take to explore parameters for the representations and how we choose the classifier for prediction of the phenotype of interest in this study

### 3.1 Body-site identification

#### (i) Bootstrapping for sampling rate selection for k-mers

Higher k values require higher sampling rates to produce self-consistent and representative representations (Figure 4). For each k, we consider a certain threshold on 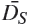 and 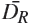 to ensure picking a sampling size resulting in self-consistent and representative representations.

**Figure 4.**
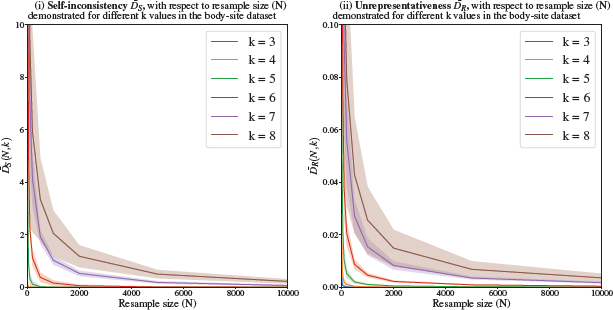
**Measuring (i) self-inconsistency 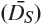, and unrepresentativeness** 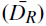 for the body-site dataset. Each point represents an average of 100 resamples belonging to 10 randomly selected 16S rRNA samples. Higher k values require higher sampling rates to produce self-consistent and representative samples.

#### (ii) Classification for different values of k with a sampling size selected based on the output of bootstrapping

Interestingly, using only 3-mer features with a very low sampling rate (≈20/15000=0.0013) provides a relatively high performance for 5-way classification. The value of macro-F1 increases with the value of k from 3 to 6, but increasing k further than that does not have any additional effect on macro-F1 (Table 1, step (ii)).

**Table 1.**
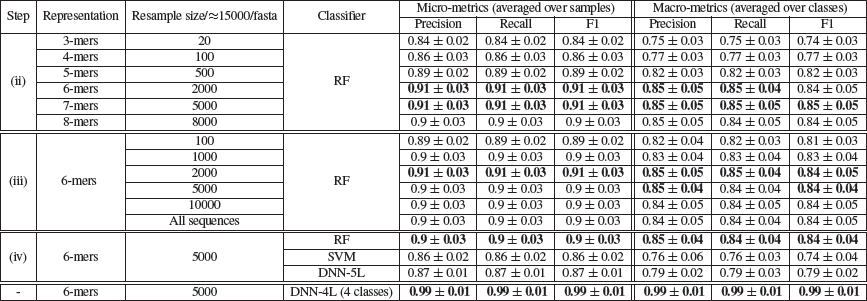
The results for classification of major body-sites using k-mer representations. The set of rows matches the steps (ii to iv) mentioned in Figure 3, i.e k-mer selection, N (sample size) selection, and finally selection of the classifier. The classifiers (Random Forest, Support Vector Machine and Neural Network classifiers) are tuned and evaluated in a stratified 10xfold cross-validation setting. The last row shows the Neural Network’s performance in the classification of body-sites when the urogenital body-sites are combined.

#### (iii) Exploring the sampling size (N) for a selected k-mer

For a selected k-value (k=6), using the Random Forest classifier for different sampling sizes are presented in Table 1, step (iii). The results suggest that changing the sampling size from 0.6% to 100% of the sequences will not change the classification results substantially, suggesting that in body-site identification, a very shallow sub-sampling of the sequences is sufficient for a reliable prediction. Using more sequences does not necessarily increase the discriminative power and may even result in over-fitting. We selected a sampling size of 5000 for 6-mers (the sampling size with the highest macro-F1 and the minimum standard deviation) for comparison between classifiers in the next step.

#### (iv) Comparison of classifiers for the se-lected N, k

In the body-site prediction task, the Random Forest classifier obtained the top macro-F1 (0.84) for this 5-way classification (Table 1, step (iv)). The confusion matrix in Figure 5 shows that the most difficult decision for the classifier is to distinguish between mid vagina and posterior fornix, both of which are urogenital body-sites. The visualizations of body-site samples obtained through using PCA, t-SNE, and t-SNE on the activation function of the last layer of the trained 5-layered Neural Network are presented in Figure 6. These results suggest that supervised training of representations using Neural Networks provides a non-linear transformation of data that can discriminate between dissimilar body-sites with reasonably accuracy. As shown in the last row of Table 1, combining the urogenital body-sites increases the macro-F1 to 0:99 0:01 using the Neural Network.

**Figure 5.**
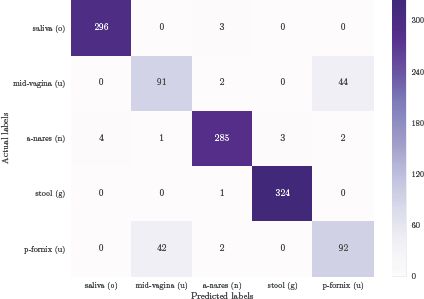
**The confusion matrix for the classification of 5 major body-sites,** using Random Forest classifier in a 10xfold cross-validation scheme. The presented body-sites are saliva (o: oral), mid-vagina (u: urogenital), anterior nares (n: nasal), stool (g: gut), and posterior fornix (u: urogenital).

**Figure 6:**
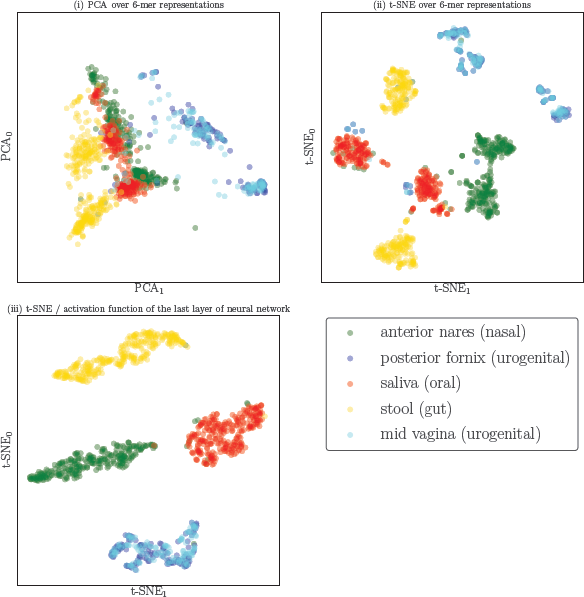
**Visualization of the body-site dataset using different projection methods:** (i) PCA over 6-mer distributions with unsupervised training, (ii) t-SNE over 6-mer distributions with unsupervised training, (iii) visualization of the activation function of the last layer of the trained Neural Network (projected to 2D using t-SNE).

#### Comparison of k-mer and OTU features in body-site identification

For a comparison between OTU features and k-mer representations in body-site identification, the Random Forest classifier (as an instance of non-linear classifier) and linear SVM (as an instance of linear classifier) were tuned and evaluated in a stratified 10xfold cross-validation setting. Our results suggest that for both k-mer features and OTUs, Random Forest is the best choice (Table 2). In addition, with almost 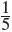 of the size of OTU features and in spite of being considerably less expensive to calculate, k-mers marginally outperforms OTU features in body-site identification.

**Table 2:**
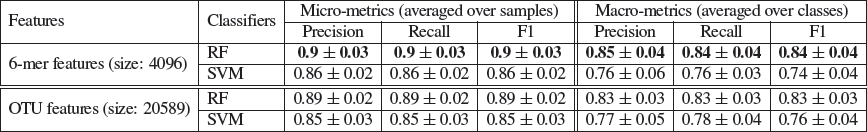
Comparison of k-mer and OTU features in body-site identification.

### 3.2 Crohn’s disease prediction

#### (i) Bootstrapping for sampling rate selection for k-mers

Similar to bootstrapping for the body-site dataset (Figure 4), 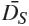 and 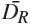 for different values of k with respect to sampling sizes were calculated. As the structure of the curve is similar to the body-site dataset, to avoid redundancy, the figure is provided as supplementary material.

#### (ii) Classification for different values of k with a sampling size selected based on the output of bootstrapping

Choosing k=6 with a sampling size of 2000 (≈2,000/38,000 = 0.05) provided a macro-F1 of 0.75 which is the minimum k with top performance (Table 3, step (ii)).

**Table 3:**
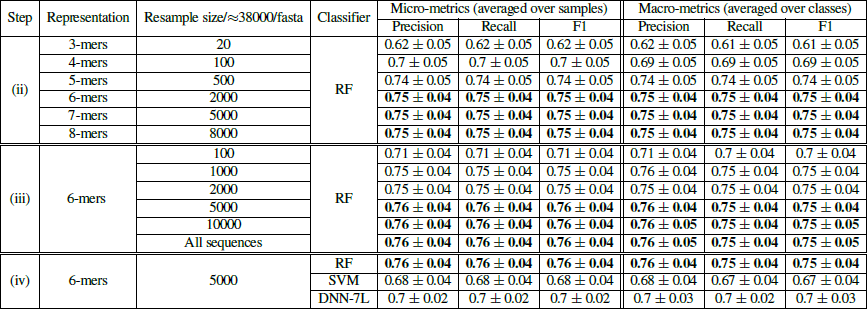
The results for classification of Crohn’s disease prediction using k-mer representations. The set of rows matches the steps (ii to iv) mentioned in Figure 3, i.e k-mer selection, N (sample size) selection, and finally selection of the classifier. The classifiers (Random Forest, Support Vector Machine and Neural Network classifiers) are tuned and evaluated in a stratified 10xfold cross-validation setting. The step column refers to the steps in Figure 3.

#### (iii) Exploring the sampling size (N) for a selected k-mer

For a selected k-value (k=6), using the Random Forest classifier for different sampling sizes while ncreasing the sampling size from 100 (100/38000=0.003) to 5000 (5000/38000=0.13) increased the macro-F1 from 0.7 to 0.75 (Table 3, step (iii)). However, using all sequences instead of 0.13 of them in each sample, did not increase the discriminative power.

#### Comparison of classifiers for the selected N, k

For selected values of k and N, the results of the Crohn’s disease prediction using Random Forest, SVM, and Neural Network classifiers are provided (Table 3, step (iv)). The Random Forest classifier obtained the top macro-F1 (0.75) for this binary classification. The projection of the Crohn’s disease dataset using with PCA and t-SNE over raw k-mer representations, as well as t-SNE over the representation learned though the supervised learning of the Neural Network are visualized in Figure 7. The trained Neural Network provides a non-linear transformation of data that can identify subjects diagnosed with Crohn’s disease with reasonable performance.

**Figure 7:**
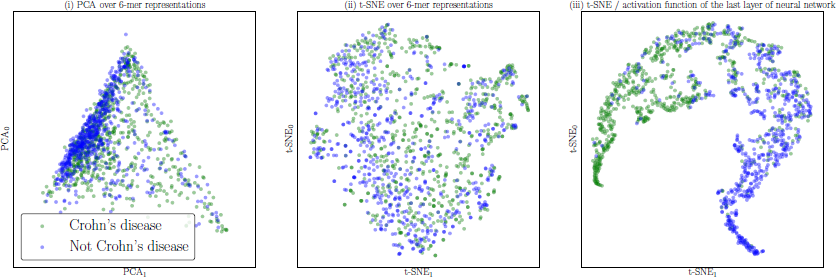
**Visualization of the Crohn’s disease dataset using different projection methods:** (i) PCA over 6-mer distributions with unsupervised training, (ii) t-SNE over 6-mer distributions with unsupervised training, (iii) visualization of the activation function of the last layer of the trained Neural Network (projected to 2D using t-SNE).

#### Comparison of k-mer and OTU features in Crohn’s disease classification

For a comparison between OTU features and our proposed k-mer representations in detecting Crohns disease from metagenomic samples, the Random Forest classifier (as an instance of non-linear classifiers) and linear SVM (as an instance of linear classifiers) were tuned and evaluated in a stratified 10xfold cross-validation. For both k-mer features and OTUs, the Random Forest classifier performed best (Table 4). In addition, even though only half of the number of features were used and in spite of being considerably less expensive to calculate, k-mers marginally outperformed OTU features in the detection of Crohns disease.

**Table 4:**
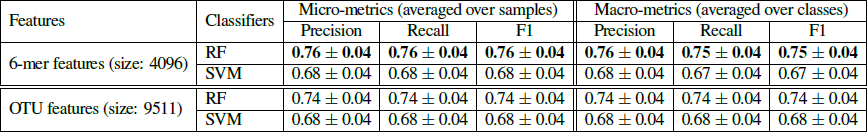
Comparison of k-mers and OTU features in the detection of the Crohn’s disease phenotype. For this comparison, Random Forest classifier (as an instance of non-linear classifiers) and linear SVM (as an instance of linear classifiers) have been used. The classifiers are tuned and evaluated in a stratified 10xfold cross-validation setting.

### 3.3 Ecological environment prediction

#### (ii) Classification for different values of k

As stated before, for the ecological and organismal datasets we do not need to perform resampling as we classify single 16S rRNA gene representative sequences. We thus can skip steps (i) and (iii) (Figure 3). Step (ii) in Table 5 shows the effect of k in the performance of the classification of the 18 environments for the ECO-18K dataset. Using k=6 provides the best classification performance, with a macro-F1 of 0.73 which is relatively high for a 18-way classification (has a mere 0.06 chance of randomly being assigned correctly for balanced dataset).

#### (iv) Comparison of classifiers for the selected k

For selected values of k, the results of the environment prediction with the Random Forest, SVM, and Neural Network classifiers are provided (Table 5, step (iv), ECO-18K dataset). SVM classifier obtained the top macro-F1 (0.79) for 18-way classification. To see the effect of increasing the number of data points in classification performance we repeat the classifier comparison (step iv) for the ECO-180K dataset. The results are summarized in Table 5-ECO-18K dataset, showing that feeding more training instances results in better training for the deep learning approach, which outperformed the SVM and achieved a macro-F1 of 0.88, which is very high for a 18-way classification task. In training the Neural Networks for the ECO-18K dataset, increasing the number of hidden layers from 3 to more did not help result in improvements. However, using the ECO-180K dataset, which is 10 times larger, allowed us to train a deeper network and increased the macro-F1 by 5 percent going from 3 layers to 5 layers. Increasing the number of layers further did not result in any improvements.

**Table 5:**
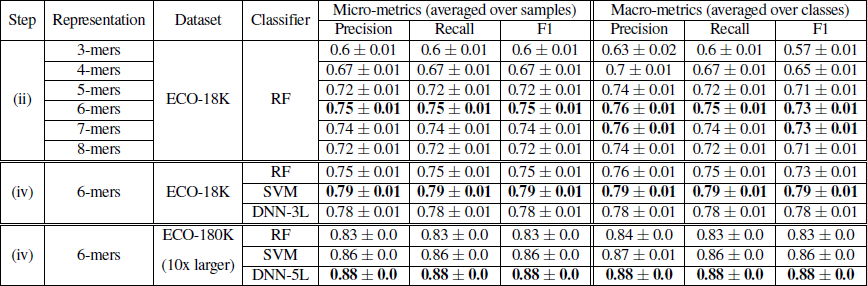
The results for the task of selecting between 18 ecological environments. The classifiers (Random Forest, Support Vector Machine and Neural Network classifiers) are tuned and evaluated in a stratified 10xfold cross-validation setting over both ECO-18K and ECO-180K datasets. The step column refers to the steps in Figure 3.

#### Neural Network Visualization

Visualizations of representative 16S rRNA gene sequences in 18 ecological environments obtained through using PCA, t-SNE, and t-SNE on the activation function of the last layer of the trained Neural Network are presented in Figure 8. For ease of visualization, we randomly picked 100 samples per class. These results suggest that supervised training of representations using Neural Networks provides a non-linear transformation of data containing information about high-level similarities between environments in the sub-plot on the right (scatter plot (iii)), where such structures appeared in the visualization only when more hidden layers were used: **(i)** On the left, the environments containing water are clustered in a dense neighborhood: marine, aquatic, freshwater, hot springs, bioreactor sludge (described in the source: “Bioreactor sludge is usually the sludge inside the bioreactor that treats waste-water.”), groundwater, and, surprisingly, rhizosphere (an environment where plants, soil, water, microorganisms, and nutrients meet and interact). **(ii)** In the middle, environments labeled related to sediment are found: sediment, freshwater sediment, marine sediment, and soil. **(iii)** Environments containing food, like food, food fermentation, and compost are at the bottom of the plot.

**Figure 8:**
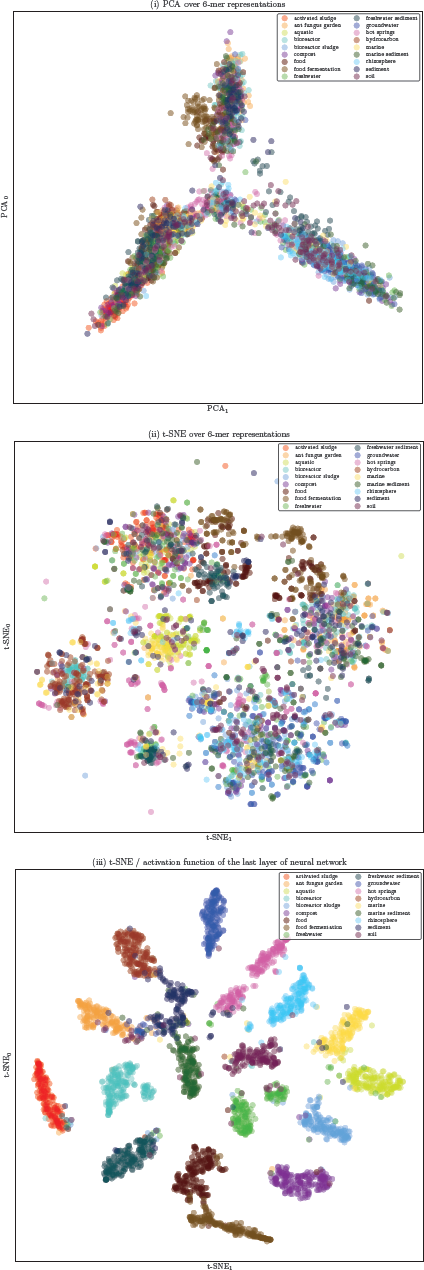
**Visualization of 18 ecologicalmicrobial environments using different projection methods:** (i) PCA over 6-mer distributions with unsupervised training, (ii) t-SNE over 6-mer distributions with unsupervised training, (iii) visualization of the activation function of the last layer of the trained Neural Network (projected to 2D using t-SNE).

#### 3.4 Organismal environment identification

##### (ii) Classification for different values of k

The results show that k=6 and 7 provide a high classifica-tion macro-F1 of 0.87 for 5 classes (0.2 chance of randomly occurring), step (ii) in Table 6.

**Table 6:**
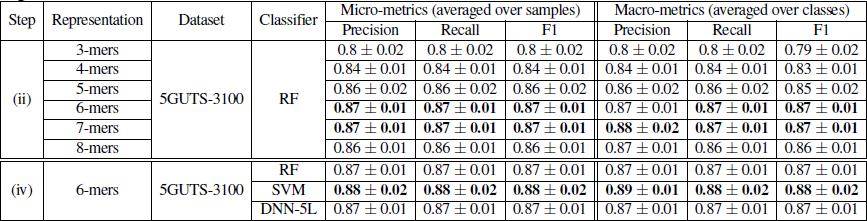
The results for the task of classifying 5 organismal environments belonging to 5 organisms’ gut. The classifiers (Random Forest, Support Vector Machine and Neural Network classifiers) are tuned and evaluated in a stratified 10xfold cross-validation setting over 5GUTS-3100 dataset. The step column refers to the steps in Figure 3.

##### (iv) Comparison of classifiers for the selected k

For selected values of k the results of the organism prediction using Random Forest, SVM, and Neural Network classifiers are provided (Table 6, step (iv)). The SVM classifier obtained the top macro-F1 (0.88) for 5-way classification.

## 4 Conclusion

In this work, MicroPheno, a new approach for predicting environments and host phenotypes on 16S rRNA gene sequencing has been presented based on k-mer representations of shallow sub-samples. We conclude with discussing the results of this study in three parts: (1) the use of k-mers versus OTUs, (2) the benefits of shallow sub-sampling, and (3) classical methods versus the deep learning approach.

### K-mers versus OTUs

To evaluate MicroPheno, we compared k-mer representations with OTU features in two tasks of body-site identification and Crohn’s disease classification. We replicated the state-of-the-art approach, i.e. Random Forest over OTU features, on datasets larger than those that were previously explored. We showed that k-mer features outperform conventional OTUs, while having several advantages over OTUs, as listed below:

- The k-mer representations are easy to compute at no computational cost for any type of alignment to references or tasks of finding pair-wise sequence similarity within samples as in OTU-picking pipelines. Just to get an idea of the computational efficiency of k-mer calculation in comparison with OTUs, note that OTU picking for the Crohn’s disease dataset of 1359 samples required more than 5 hours using 5 threads, while 6-mer distribution calculation took around 5 minutes using the optimal sampling size (N) for classification using the same number of threads.
- Taxonomy-independent analysis is often the preferred approach for amplicon sequencing when the samples contains unknown taxa. k-mer features can be used without making any assumptions about the taxonomy. However, OTU-picking pipelines make assumptions about the taxonomy as discussed in §1.2; therefore they can even be phylogenetically incoherent.
- k-mer distribution is a well-defined representation, while OTUs are sensitive to the pipeline and the parameters.
- Sequence similarities are naturally incorporated in the k-mer representations for the downstream learning algorithm, while with grouping sequences into certain categories, the sequence similarities between OTUs are ignored.

The main disadvantage of k-mer features over OTUs is that using short k-mers makes it more difficult to trace the relevant taxa to the phenotype of interest. When such an analysis is needed, using OTUs or increasing the size of k would be an alternative solution. However, as long as prediction is concerned, using a k-mer representation seems to be the best choice for an accurate and rapid detection/diagnosis over 16S rRNA sequencing samples.

### Shallow sub-sampling

We proposed a bootstrapping framework to investigate the consistency and representativeness of k-mer distributions for different sampling rates. Our results suggest that, depending on the k-mer size, even very low sub-sampling rates (0.001 to 0.1) (for k between 3 to 7) not only can provide a consistent representation, but can also result in better predictions while possibly avoiding overfitting. Setting aside the save in preprocessing time as a natural benefit of sampling, this result also suggest that at least for similar phenotypes of interest, shallow sequencing of the microbial community would be sufficient for an accurate prediction.

### Classical classifiers versus deep learning

We studied the use of deep learning methods as well as classic machine learning approaches for distinguishing among human body-sites, diagnosis of Crohn’s disease, and predicting the environments from representative 16S gene sequences. Studying the role of dataset size in the classification of ecological environments showed that for large datasets using deep learning provides us with more accurate predictions. However, when the number of samples are not large enough, Random Forests performs better on both OTUs and k-mer features. In addition, we observed that for classification over representative sequences as opposed to samples (pool of sequences), the SVM outperformed the Random Forest classifier. Another advantage of using deep learning in classification was that supervised training of a proper representation of data results in a more discriminative representation for the downstream visualization compared to the unsupervised methods (PCA and t-SNE on the raw k-mer distributions). For body-site identification and even more clearly in ecological environment classification, the model was able to extract more high-level similarities between the environments.

